# Recovery Dynamics of the High Frequency Alternating Current Nerve Block

**DOI:** 10.1101/235135

**Authors:** Adrien Rapeaux, Emma Brunton, Kianoush Nazarpour, Timothy G. Constandinou

## Abstract

**Objective:** High-Frequency alternating current (HFAC) nerve block has great potential for neuromodulation-based therapies. However nerve function recovery dynamics after a block is highly understudied. This study aims to characterise the recovery dynamics of neural function after an HFAC block.

**Approach:** Experiments were carried out *in-vivo* to determine blocking efficacy as a function of blocking signal amplitude and frequency, and recovery times as well as recovery completeness was measured within a 0.7 s time scale from the end of block. The sciatic nerve was stimulated at 100 Hz during recovery to reduce error to within ±10 ms for measurements of recovery dynamics. The electromyogram (EMG) signals were measured from *gastrocnemius medialis* and *tibialis anterior* during trials as an indicator for nerve function.

**Main Results:** The HFAC block was most reliable around 20 kHz, with block thresholds approximately 5 or 6 mA depending on the animal and muscle. Recovery times ranged from 20 to 430 milliseconds and final values spanned relative outputs from approximately 1 to 0.2. Higher blocking signal frequencies and amplitudes increased recovery time and decreased recovery completeness.

**Significance:** These results confirm that recovery dynamics from block depend on blocking signal frequency and amplitude, which is of particular importance for neuromodulation therapies and for comparing results across studies using different blocking signal parameters.

## 1. Introduction

Recent developments in neurotechnology have established new modalities for interacting with the body’s nervous system. The visions of phrama- and electroceutical companies such as GSK and Verily’s Galvani Bioelectronics [1] and Feinstein Institute [2], as well as other governmental programmes run by DARPA, e.g. SPARC and ElectRx [3], is to create highly miniaturised and powerful implants targeting the body’s nervous system for therapeutic effect. This has brought about a change of paradigm where the goal of stimulation is not only to excite nerve cells but also to inhibit them, paving the way for novel dynamic neuromodulation therapies. An essential tool in this endeavour is High Frequency Alternating Current (HFAC) nerve block, which is under active research [4, 5, 6, 7, 8].

The HFAC nerve block is a powerful stimulation technique that allows complete and reversible inhibition of action potential conduction in the peripheral motor and autonomous nervous system [9, 10]. It has proved effective in several species including the rat [11], frog [12] and non-human primates [13], amongst others. In addition, blocking neural transmission via the vagus nerve with HFAC has been used clinically for the treatment of obesity [9, 10]. Other applications include bladder control [14, 15] and pain management [6].

Some main characteristics of the HFAC nerve block are understood experimentally. These include its quick onset [16] and recovery [4] and the intense transient neural firing observed at initial application of the blocking signal, that is termed the onset response. However many aspects of the HFAC nerve block are still understudied. For instance, characteristic phenomena, such as the onset response, have not yet been successfully modeled *in silico*, motivating experimental investigation of block. The exact mechanism of action of block remains uncertain as researchers have disagreed on whether block is caused by sodium-related depolarisation of the membrane [17] or potassium-related effects [18]. Models of HFAC block today do not capture all the phenomena that are reported in experimental literature such as how different fibre types react differently to block [19], or how applying block for different durations can change the rate at which the nerve recovers to initial excitability [20, 7].

Understanding the recovery dynamics of the HFAC nerve block can yield improvements in selectivity or power consumption for stimulators, for example by exploiting recovery dynamics to reduce the duty cycle of blocking [21]. While current studies are investigating recovery dynamics from block applied over long periods of 15 minutes of more [7], relatively little attention has been brought to investigating recovery of block on shorter timescales, where recovery is almost instantaneous. The time needed for the nerve to recover from block when the signal has been applied for less than 10 minutes is typically in the order of one second [12], however no precise measurements have been made of this duration.

A precise measurement of the recovery time for block would enable reductions in power consumption for stimulators by use of duty-cycling, and if different nerve fibres recover at different speeds the resulting dynamics can be exploited for selective stimulation, reducing side-effects for therapies such as vagus nerve stimulation [22] or bladder stimulation [23].

In this study, we set out to characterise the recovery dynamics of an HFAC block in the rat sciatic nerve. We used two separate muscular activity outputs measured using the EMG signal to quantify the recovery dynamics. The protocol and methodology are explained in detail, wherein rats were implanted with a custom cuff to carry out simultaneous block and stimulation to measure the muscle’s response as it was recovering from block.

## 2. Materials and Methods

### 2.1. Ethics and Veterinary Surgery

All animal care and procedures were performed under appropriate licences issued by the UK Home office under the Animals (Scientific Procedures) Act (1986) and were approved by the Animal Welfare and Ethical Review Board of Newcastle University.

Four Sprague-Dawley rats weighing 350-450 grams were used in this study. General surgery procedures followed those described in [24]. Briefly, animals were initially anaesthetised by either an intraperitoneal injection of medazolam and fentanyl/fluanisone (hypnorm) with an initial dose of 2.7mL/kg [25], or in a box with 3% isoflurane in oxygen. Anesthesia was maintained by isoflurane in oxygen (0.5 to 3%, adjusted as required) through a nose cone and adequate anesthetic depth was indicated by absence of withdrawal to a noxious toe pinch, and the presence of a regular heart and respiratory rhythm. Temperature, pulse and breathing rates were monitored throughout the procedure.

Figure 1a depicts the experimental setup. After shaving and cleaning, an incision was made on the dorsal aspect of the right leg to expose muscle covering the sciatic nerve. The muscles were blunt-dissected with surgical scissors to expose the sciatic nerve. A custom-built split nerve cuff (Microprobes) was implanted on the exposed nerve length with the slit facing upwards. Cuff contacts were made from 0.1 mm diameter platinum wire. The inter-contact distance was 0.5 mm for the two extremity tripoles and 1mm for the 8-contact middle ladder. The distance separating the tripoles from the ladder was 3 mm. The silastic material overhanged the extremity tripoles by 1 mm. The total length of the cuff was 24 mm. The embedded sutures were used to close the cuff around the nerve, and Kwik-Cast (World Precision Instruments) silicone elastomer was applied to further secure and isolate the cuff and nerve. The *gastrocnemius medialis* and *tibialis anterior* muscles were exposed by incision of the skin and implanted with separate pairs of insulated tungsten wires (Thickness: 100 *μ*m, Advent Research Material). The spine was exposed by incision and blunt dissection and an additional tungsten wire was wound around an exposed spinous process and secured with surgical cement to provide an electric ground point for the EMG amplifier. All incisions and muscles were sutured closed and back together where appropriate to minimise movement of the cuff and wires during stimulation and to prevent the tissue from drying.

**Figure 1.**
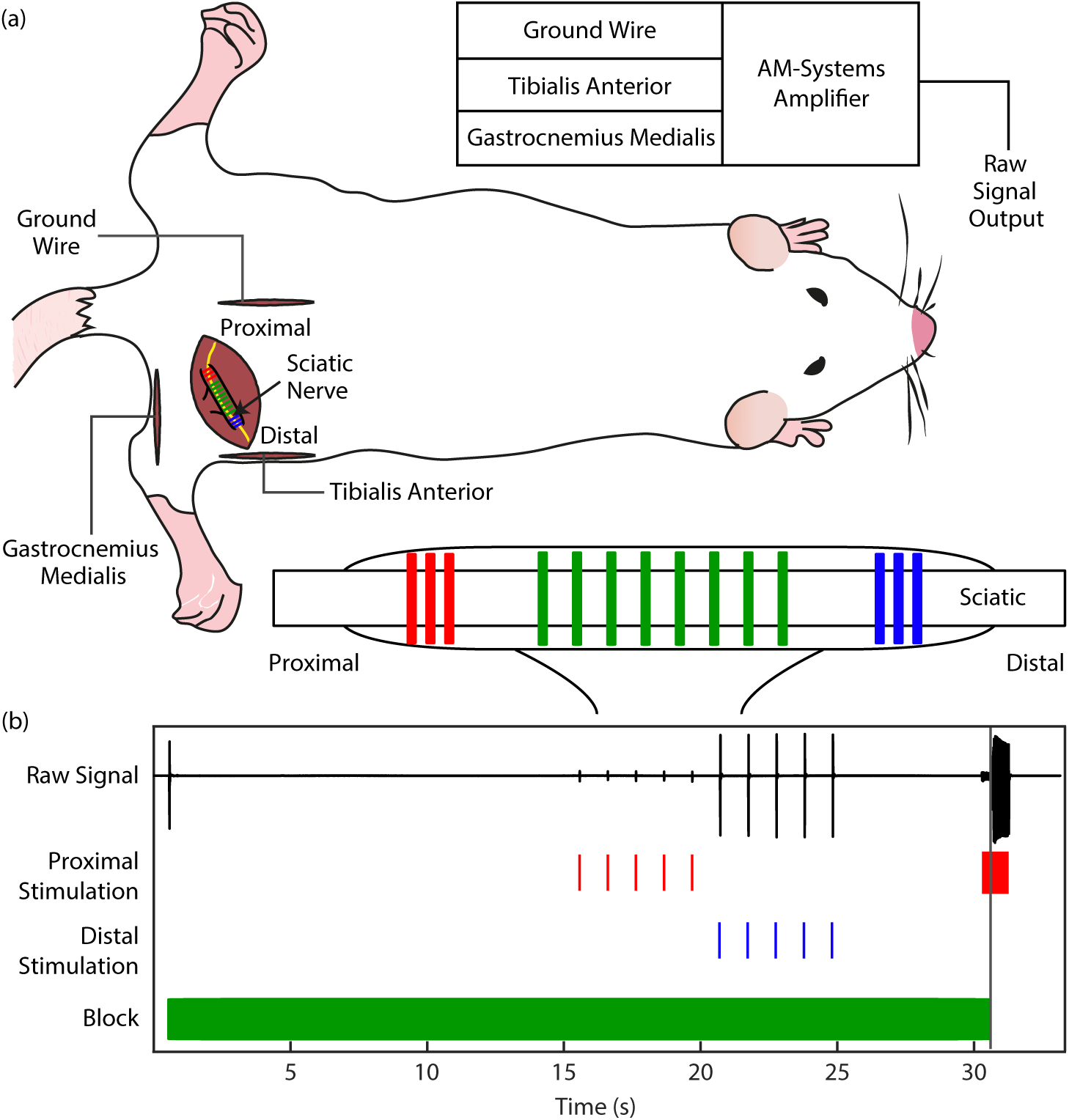
(a) Experimental setup illustrating a Sprague Dawley rat, the custom cuff electrode used and connections to the recording amplifier for the experiment. Connections to stimulating instruments not shown. Conventional stimulation was carried out by the Cerestim on cuff contacts 1-3 and 12-14. Block was carried out on cuff contacts 4-11. The Cerestim shared the ground reference with the A-M Systems amplifier. (b) Typical stimulation and block timeline showing raw EMG trace, proximal, distal and blocking stimuli. The effect of block on the EMG trace is visible as proximally elicited activity is subdued as long as block is active.

### 2.2. Devices and Setup

The tungsten wires used to capture EMG signals from *gastrocnemius medialis* and *tibialis anterior* were connected to two channels of a differential amplifier (Model 1700, A-M Systems, WA, USA) as shown in Figure 1a. The EMG signals were filtered using a 10 Hz highpass second-order filter, a 10 kHz lowpass second-order filter, and a 50 Hz notch filter to remove line noise. The amplifier gain was set to 100. The output of the differential amplifier was connected to two analogue channels of a Cerebus Neural Signal Processor (Blackrock Microsystems, Utah, USA).

The cuff contacts were connected to two different devices. The three contacts on each end were connected to a Cerestim stimulator (Blackrock Microsystems, Utah, USA) while the middle eight contacts were connected to a custom-built current-controlled HFAC stimulator printed circuit board (PCB). The CereStim system carried out stimulation distal and proximal to the area blocked by HFAC. The *sync* signal from the CereStim was connected to one of the analogue channels of the CereBus, so that the EMG signals could be synchronised with the stimulation.

The schematic for the analogue portion of the circuit is shown Figure 2, with component values detailed in Table 1.

**Figure 2.**
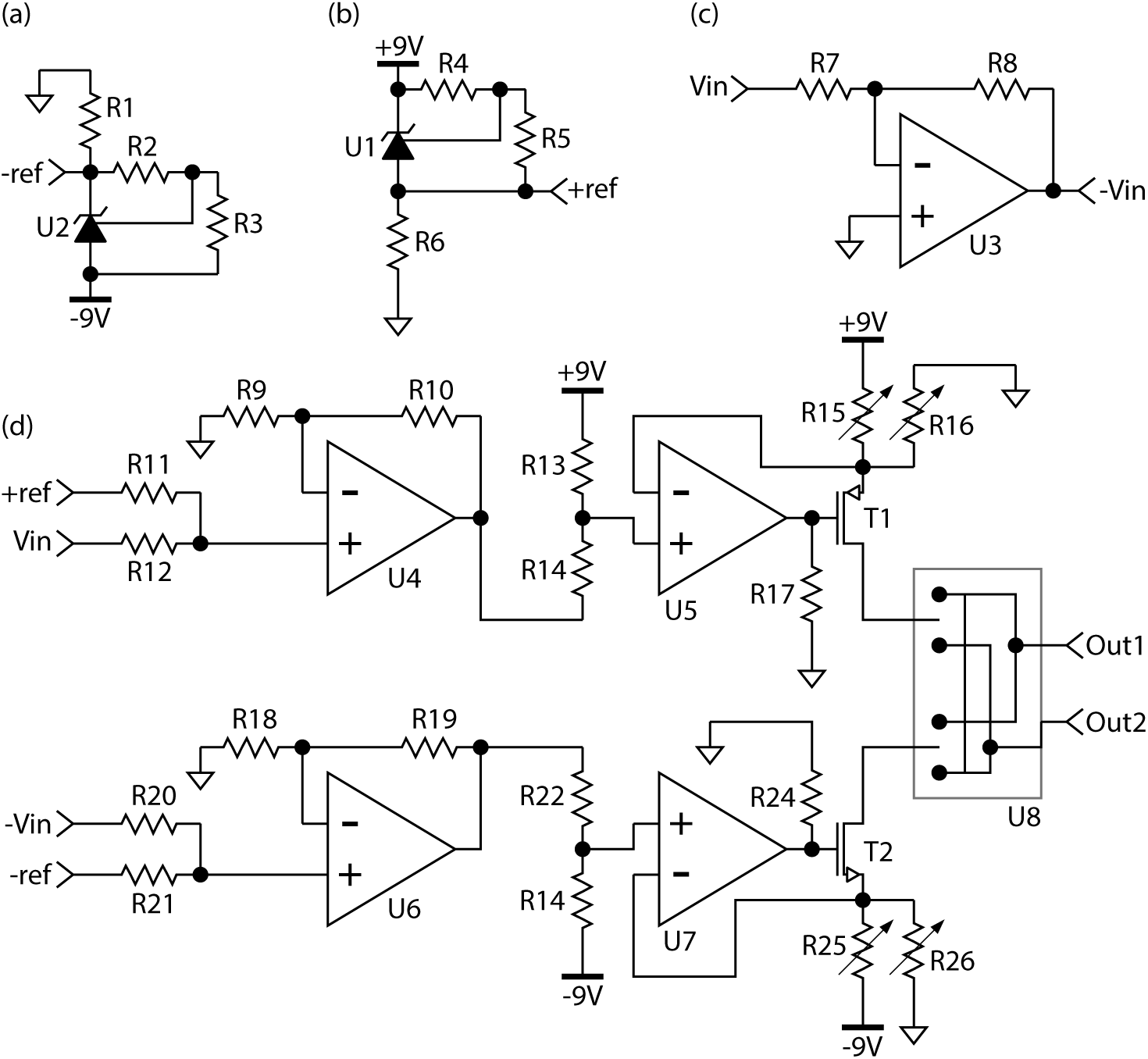
Circuit schematic for the analogue portion of the custom-made PCB device used to deliver the blocking signal. (a) and (b) are voltage references built using precision shunts; (c) is an inverter used to invert the DAC-generated input signal to the opposite polarity for input into the sink branch of the stimulator; and (d) is the core of the stimulator with separate current source and sink individually calibrated with trim potentiometers. Connections between parts of the circuits are named using node references.

**Table 1.** Component values and designators for Figure 2. The symbol R denotes resistor. The sign Ω, that is Ohms, is dropped for brevity.

| Part | Value | Part | Value | Part | Value | Part | Value |
| --- | --- | --- | --- | --- | --- | --- | --- |
| R1 | 5 k | R10 | 10 k | R19 | 10 k | R26 | 20 k |
| R2 | 10 k | R11 | 10 k | R20 | 10 k | U1 | LM4041 |
| R3 | 14.7 k | R12 | 10 k | R21 | 10 k | U2 | LM4041 |
| R4 | 10 k | R13 | 1 k | R22 | 27 k | U3 | OPA192 |
| R5 | 15.4 k | R14 | 27 k | R23 | 1 k | U4 | OPA4192(a) |
| R6 | 5 k | R15 | 50 | R24 | 50 k | U5 | OPA4192(b) |
| R7 | 50 k | R16 | 20 k | R25 | 50 | U6 | OPA4192(c) |
| R8 | 50 k | R17 | 50 k | T1 | BSS8402(a) | U7 | OPA4192(d) |
| R9 | 10 k | R18 | 10 k | T2 | BSS8402(b) | U8 | ADG1433 |

The custom PCB device drove the high frequency blocking signal through blocking capacitors to prevent DC contamination. Switching and TTL signals were driven by a KL25z Microcontroller (Freescale) to ensure consistent timing in trials, and a MATLAB^™^ (version: 16a) graphical user interface was used to transfer blocking signal parameters to the device. Both current branches were calibrated by adjustment of the trim potentiometers shown to output the same current with opposite polarity, with a common 12-bit digital to analouge converter input. Devices were powered by galvanically-isolated power supplies.

### 2.3. Experimental Protocol

#### 2.3.1. Electrode Impedance Measurement

Upon implantation of the nerve cuff, contact impedance relative to the grounding wire was measured with the Cerebus to evaluate contact quality. Contact quality was considered acceptable when impedances were below or around 10 kΩ at 1kHz.

#### 2.3.2. Supramaximal Stimulation Threshold Determination

A standard biphasic cathodic first stimulation waveform was used to test connection quality between each contact and the nerve. The cathodic and anodic phases are 350 *μ*s in duration, with 100 *μ*s of interphase, cathodic and anodic amplitude identical. Stimulation amplitude was progressively increased until corresponding EMG activity reached a plateau to ensure recruitment of as many nerve fibres as possible. This stimulation amplitude was then used for the remaining trials in the experiment.

#### 2.3.3. Determining the Best Block Electrode Pair

Block was tested for each adjacent electrode pair along the 8-contact middle ladder, with a standardised test at 40 kHz, 6 mA amplitude. Block at this amplitude and frequency is generally reliable, with a linear relationship between signal amplitude and effect. Block quality was determined by visual inspection of onset and twitch response due to blocking and stimulation, respectively. A lower proximal stimulation response with a short onset response and consistent distal response indicated high quality block. The electrode pair with the best block quality was chosen as the blocking pair for the rest of the trials in the experiment.

#### 2.3.4. Block Trials

For each block frequency from 10 kHz to 50 kHz in 5kHz steps, a sweep of blocking amplitudes was carried out from 2 mA to 9 mA in steps of 1mA, except for rat 1 for which the highest tested amplitude was 8 mA. In each trial, block was turned on for 30 seconds and an onset response was observed. Five proximal pulses followed by 5 distal pulses were delivered to measure block efficacy for both muscles, at a rate of 1 Hz, 15 seconds after block was turned on. Proximal pulses were used to measure block efficacy and distal pulses were used to ensure the neuromuscular junction wasnt fatigued and that the muscle responded identically to baseline as an in-trial positive control. Approximately 0.3 seconds before block cut-off, a train of stimulation pulses were delivered proximally for 1 second, at a rate of 100 Hz, to measure recovery of the nerve from block as it is turned off (Figure 1). A timeline with an example recording is shown Figure 1, for a blocking signal frequency and amplitude of 15 kHz and 6 mA, respectively. To prevent drifting baseline bias over the course of the experiment, at regular intervals the baseline excitability of the neuromuscular system was recorded with a standard distal and proximal stimulation test with no block, against which muscle activity in trials was measured.

### 2.4. Data Analysis

To obtain an objective measure of the HFAC block efficacy for one frequency-amplitude pair, the EMG activity resulting from proximal stimulation during block was compared to baseline activity without, all other variables unchanged. During recording, a digital synchronisation signal indicates stimulation times in each recording. As shown in Figure 3, this was used to extract 10 millisecond snippets from the original signal using a rectangular window containing the resulting EMG spike from stimulation. Stimulation artefacts were removed by removing the first 2 milliseconds of signal from further analysis (zeroing in visualization). The average value of the signal was then removed to avoid baseline drift bias. The absolute value of the signal was calculated and then integrated to give a value for the EMG spike. The values of the 5 spikes in each trial are averaged to give a final value. These values were normalised with respect to each experiments baseline, determined by measuring the spike values without block. For measuring the dynamics of recovery from block, the EMG spike values are measured in the 100Hz stimulation train using the aforementioned protocol then are fitted to a sigmoid curve with three regions corresponding to the lowest 10% of values, the middle 80% and the highest 10%, with examples for *tibialis anterior* and *gastrocnemius medialis* shown Figure 4. As muscle recovery changes significantly with experimental parameters and recovery is only observed when a good quality, stable block has been previously established, only the fits with an adjusted R-square of more than 0.9 with respect to the raw recovery curve were plotted. This ensured that the data corresponds to good quality block trials. regardless of whether block was complete or partial. The time between block cut-off and the end of the second (blue) region was measured as the time to recovery in seconds and the average value of the third region was measured as the final recovery value during the trial. Note that for *gastrocnemius medialis*, due to the rapid onset of fatigue as can be seen during a 100Hz frequency stimulation train without block on Figure 5, the maximum value of the measured recovery curve was used to fit for the maximum value of the sigmoid curve and thereby reduce biasing of the result by early onset fatigue.

**Figure 3.**
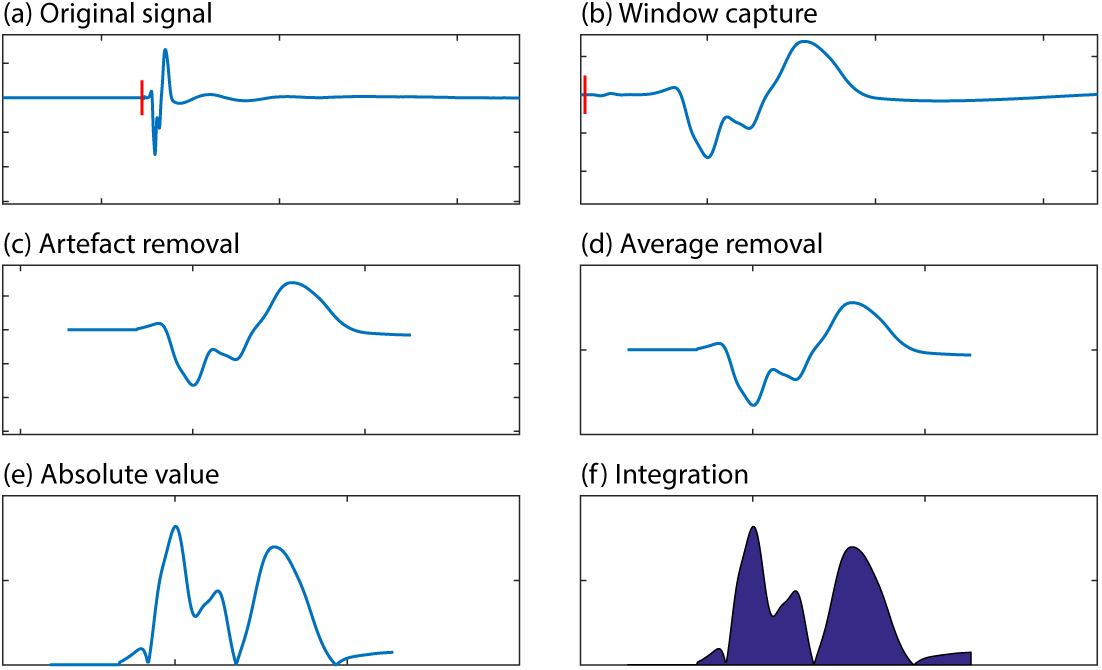
Graphs showing data; analysis process for a single EMG spike, sequentially from (a) to (f): (a) original signal before any processing; (b) a 10 ms snippet is extracted using the stimulation signal marker (red); (c) stimulation artefact is removed by not considering the first 2 milliseconds of waveform in further processing (zeroing in visualization); (d) removal of the mean of the signal; (e): calculated absolute value; (f) integration of the signal to yield a spike value.

**Figure 4.**
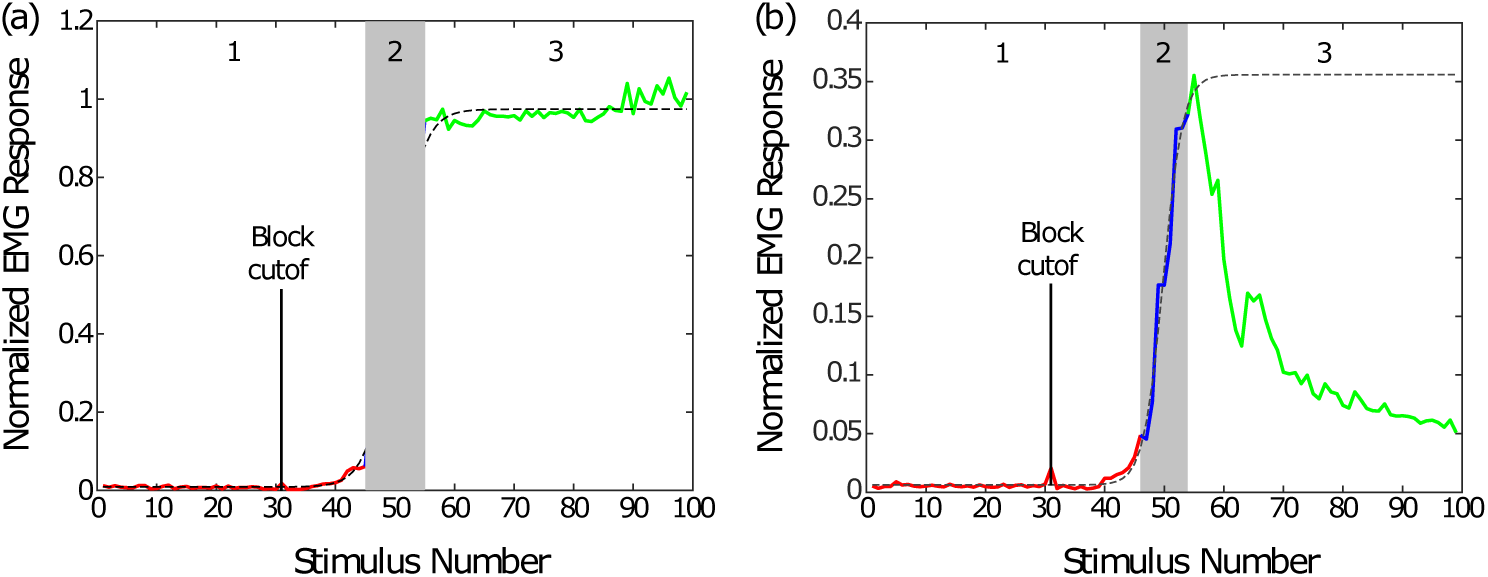
(a) Plot showing the sigmoid fit curve for *tibialis anterior* at block signal frequency 20 kHz and amplitude 7mA.(b) Plot showing the sigmoid fit curve for *gastrocnemius medialis* at block signal frequency 25 kHz and amplitude 7 mA. The three numbered regions are marked using numbers and grayscale overlay. Note the different scales in both plots, corresponding to different degrees of recovery for each muscle for different frequencies.

**Figure 5.**
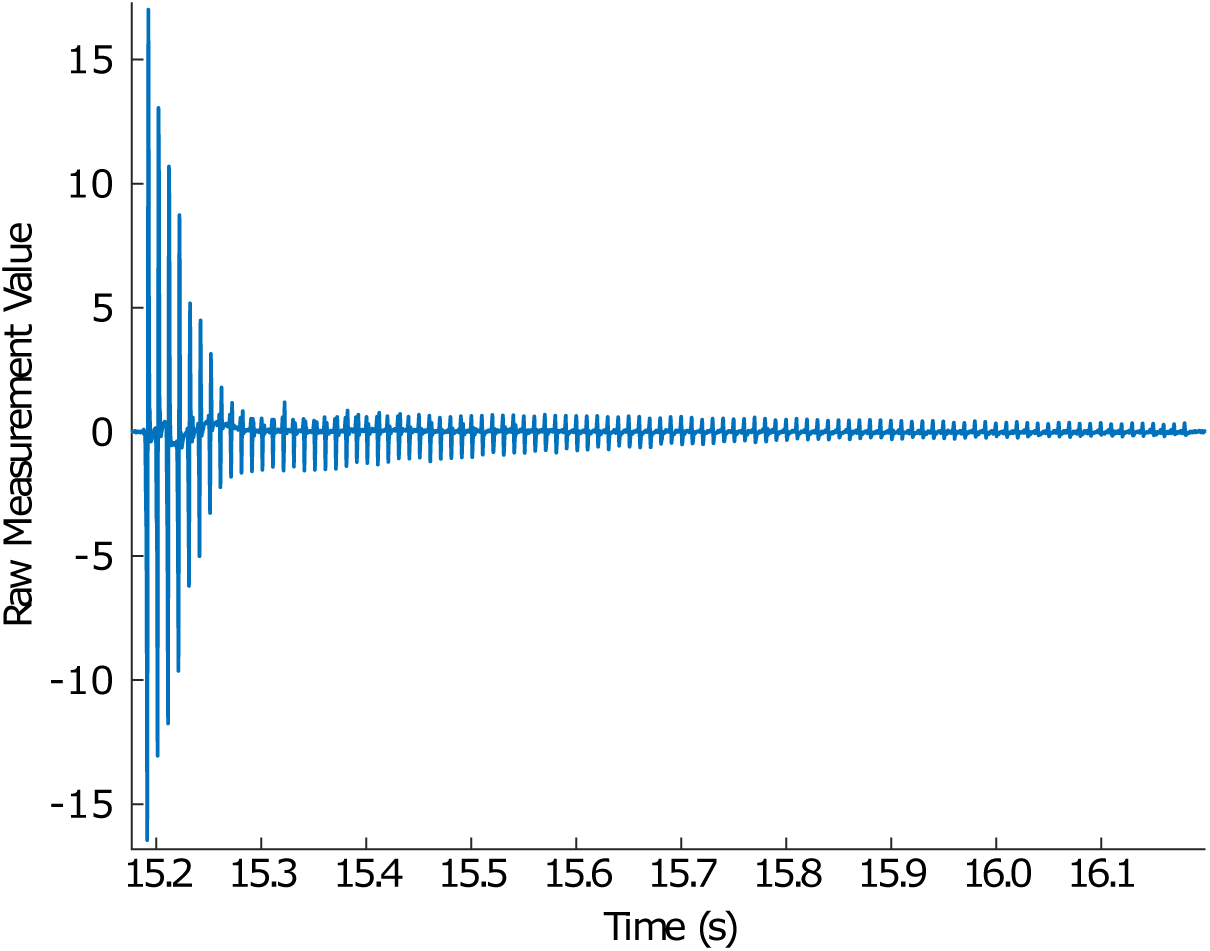
Plot of raw data during a 100Hz stimulation train carried out without block for *gastrocnemius medialis*, showing the quick onset of fatigue.

The boxplots collate data from all frequencies for which the block reduced normalised EMG activity by at least 0.5 relative to baseline that also featured sigmoid fits for which the adjusted R-square value was higher than 0.9.

## 3. Results

Several notable trends are visible in the results for block efficacy over frequency and amplitude in both muscles, plotted Figures 6 and 7. For a given frequency, an increase in the amplitude of the blocking current yields stronger effects and the resulting muscular activity from stimulation during block falls. The lowest values indicating complete block are obtained for lower frequencies, while for higher frequencies higher spike values and therefore partial blocks are obtained. It is of note that muscle response is more variable for the lowest frequencies such as 10 and 15 kHz than it is for higher frequencies, suggesting that the most reliable frequencies for block are in the 20 to 30 kHz range, where values are the most stable while maintaining strong block effects at amplitudes as low as 6 mA. A second observation is that block of *tibialis anterior* was more reliable and stable than for *gastrocnemius medialis* as seen from the lower variance of results, especially for lower block frequencies. Finally, in two rats the blocking effect was progressively lost as the blocking signal amplitude was increased past 5 mA at the lowest frequencies. Only partial block was obtained for blocking signal frequencies over 35 kHz. As previously reported in the literature this suggests there are optimal frequencies for block where lower amplitudes and frequencies achieve a strong blocking effect [12, 26].

**Figure 6.**
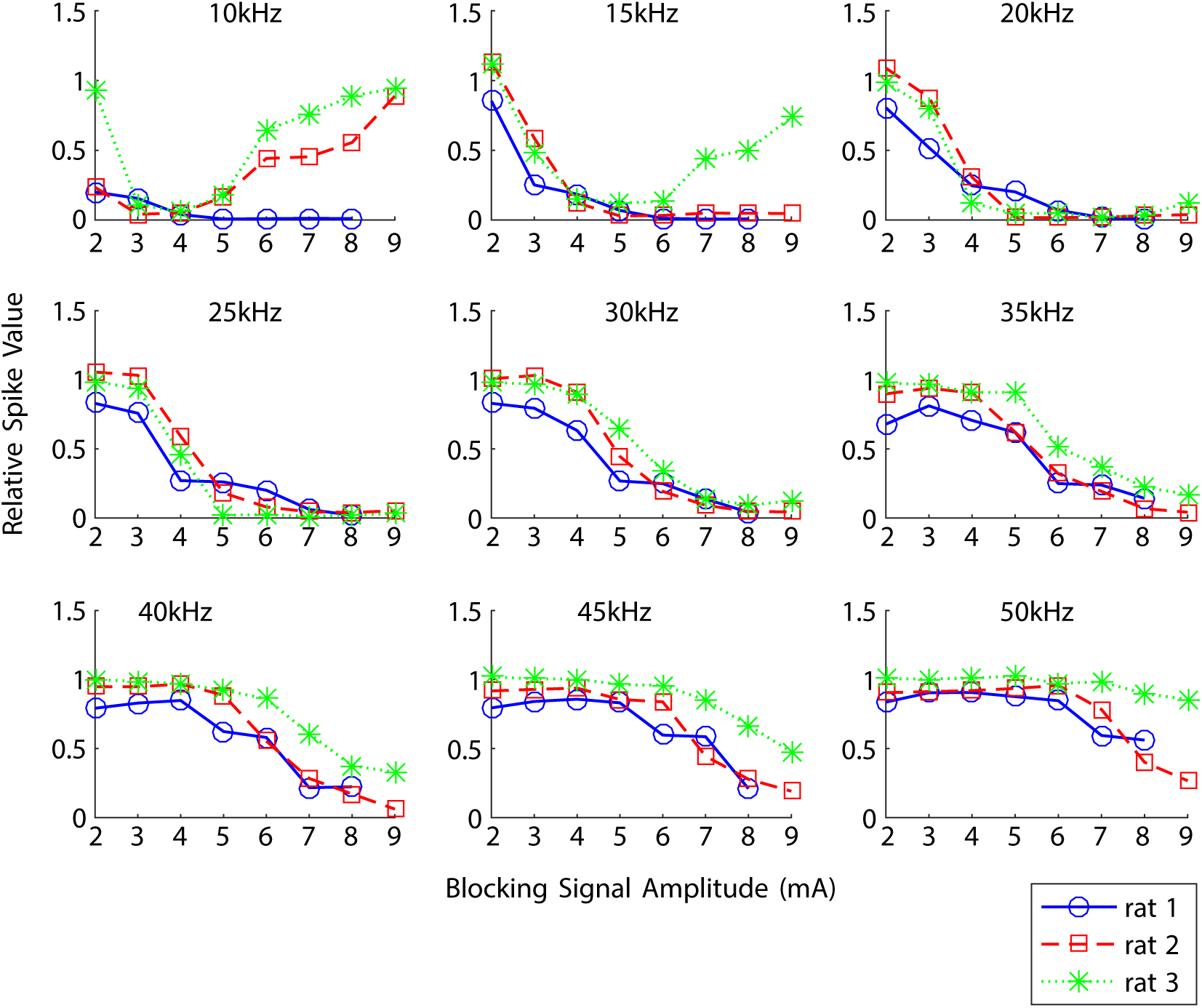
Spike value relative to baseline during block for *tibialis anterior* versus blocking signal amplitude. Each plot corresponds to a blocking signal frequency.

**Figure 7.**
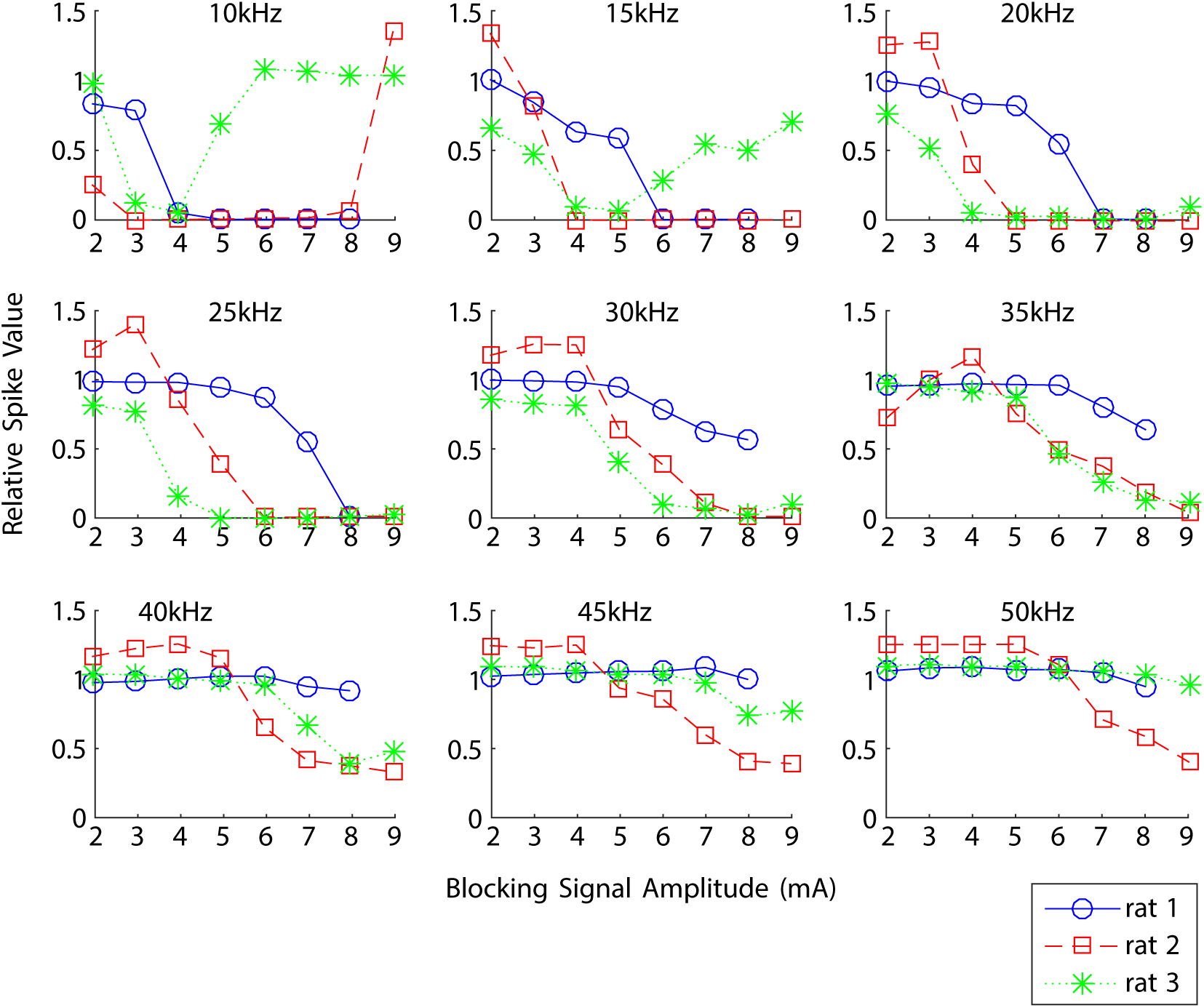
Spike value relative to baseline during block for *gastrocnemius medialis* versus blocking signal amplitude. Each plot corresponds to a blocking signal frequency.

### 3.1. Block Recovery

Block recovery data for time to recovery and final normalised EMG response are shown Figure 8, (a) and (b) respectively. Note that recovery delay values close to 0.7 indicate that the muscle activity did not plateau during the 0.7 seconds after the block was cut off (Figure 8), and that very low (under 0.25) final normalised EMG responses (Figure 8) indicate poor recovery, however subsequent controls after those trials showed no difference in muscle activity not under block. Rough trends are visible when considering average values of the data, such as an increase in recovery time and a decrease in the final normalised EMG value with increasing block amplitude. When good quality block was established, recovery times varied from around 20 milliseconds to 430 with high variance from experiment to experiment, however in the vast majority of cases recovery occurs between 100 to 300 milliseconds. For final normalized EMG responses variation in results from experiment to experiment is more pronounced, with trials from rat 3 showing much higher final recovery values than for rats 1 and 2. Final normalised EMG responses tended to be lower at higher block amplitudes for rats 1 and 2, whereas this parameter was largely insensitive to block signal amplitude for rat 3, but decreased with increasing block signal frequency (not shown).

**Figure 8.**
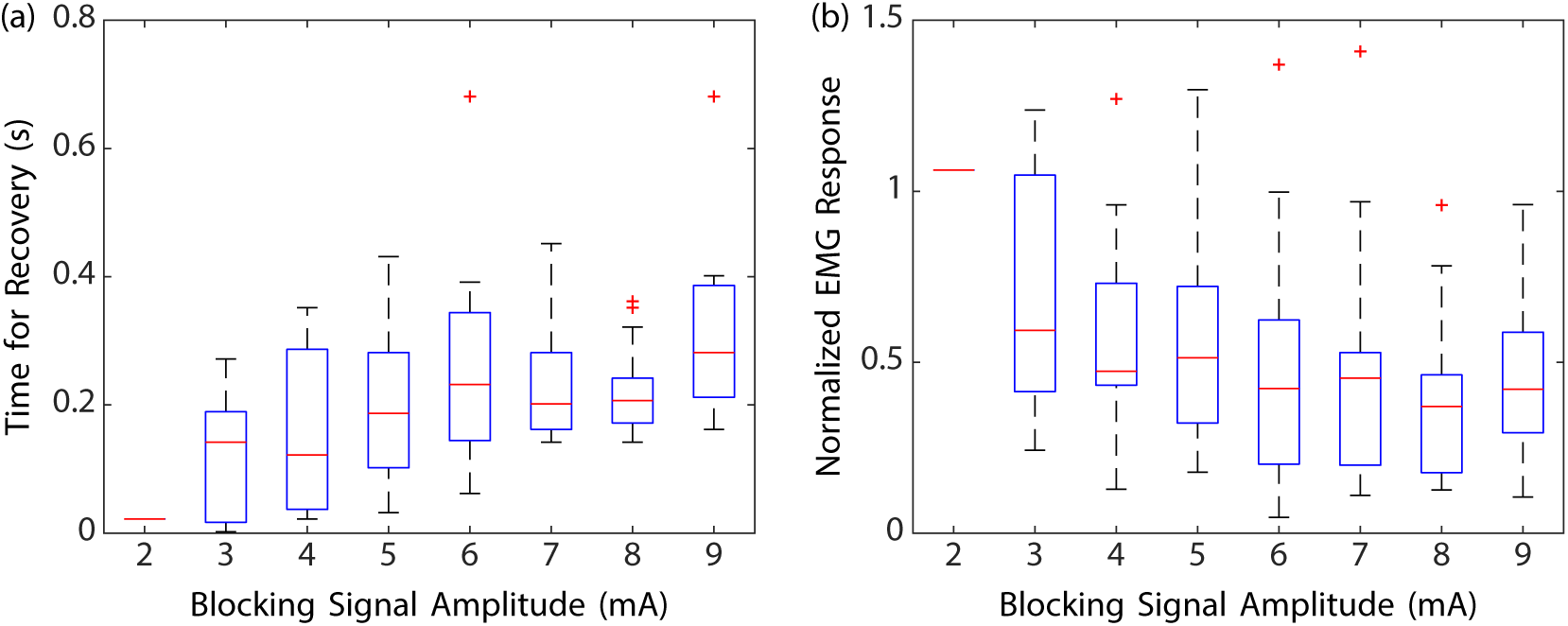
(a) Boxplots of block recovery delay in seconds versus blocking signal amplitude for all frequencies and both muscles; (b) Boxplots of final normalised EMG response at the end of the 100 Hz stimulation train versus blocking signal amplitude for all frequencies and both muscles.

## 4. Discussion

Several aspects of the results warrant discussion as variability is evident from the curves and boxplots. Measurements displayed more variance for *gastrocnemius medialis* compared to *tibialis anterior* across all three rats. It is not certain whether this is due to the difference in fiber composition innervating the two muscles or to a particular aspect of the protocol. While blocking signals at 10 kHz result in block for some amplitudes, and it is possible that block could have been obtained at lower frequencies, the results suggest that block obtained at those frequencies is unreliable from experiment to experiment, with differences in block efficacy at the same amplitude and frequency. The blocking signal frequencies yielding the most consistent results are in the 20 to 30 kHz range, where block was obtained at roughly the same amplitudes for all three rats. However, establishing statistical significance for any trend is not possible with the data obtained.

Several sources of variability were identified. Some is contributed by collation of data from both muscles. This is justified by the sparsity of the data due to the filtering detailed in subsection 2.4. In addition, for *gastrocnemius medialis* the fitting algorithm applied to the recovery curve was made to ignore data points beyond the highest relative EMG value in order to avoid fatigue bias, as rapid onset of fatigue was evident in stimulation-only trials. This fatigue could have been avoided by reducing the frequency of the end-of-trial stimulation train, however this would have been at the cost of time resolution to determine the recovery time, as performing more trials per data point would have made the experiment impractically long. Care was taken to allow sufficient time in between trials for the muscle to recover after high-frequency stimulation, coupled with periodic monitoring of baseline muscle activity to provide an adaptive baseline measure, however despite this it is possible that *gastrocnemius medialis* did not recover fully within the trials due to fatigue occurring too quickly.

Furthermore, some variance in the results may be due to the incremental changes made to the protocol at the time of the experiments, as this was still under development. The measures in Rat 3 in particular were more consistent than in the first two rats, suggesting that further work with the refined protocol would improve consistency of results and provide clearer trends. The most notable improvements to the surgical protocol were the use of external loops of suturing thread to close the cuff around the nerve rather than using the embedded sutures (see Figure 1). Kwik-Cast was applied to stabilise the assembly resulting from this setup. An added benefit is that the cuff can be recovered without damage since the loops can be simply cut.

Finally, in some trials relative EMG response decreased sharply immediately after cessation of block. At the time the experiments were carried out the reason for this was unknown, however a subsequent literature search revealed that DC discharge due to charge accumulation on DC blocking capacitors may explain these observations [27]. The DC pulses resulting from charge accumulation on the blocking capacitors in HFAC block scenarios may have lengthened block recovery times and reduced the final normalized EMG response.

Nevertheless, the results display trends that can be tentatively explained. The effect of blocking signal amplitude can be explained by reasoning that an increase in amplitude increases the charge delivered per cycle to the nerve. This leads to a stronger blocking effect which explains why the muscle requires more time to recover from the blocked state, and supports dose-dependent blocking effects [7]. This translates to the longer recovery time and lower normalised EMG response at the end of the trial. Frequency is expected to affect blocking in the sense that more charge is delivered per cycle at lower frequencies, therefore resulting in stronger block. This explains why lower frequencies reach blocking threshold at lower amplitudes, an observation that is supported in the literature [12, 15]. DC pulsing effects may have affected the results for the lowest frequencies, which could explain the variance seen, as it is expected that DC pulsing is stronger with increasing block amplitude and decreasing frequency. Further work could attempt to more effectively separate the effects of the two variables by normalising results with respect to injected charge per cycle of the blocking signal.

## 5. Conclusions

In this study the recovery from HFAC block of the *gastrocnemius medialis* and *tibialis anterior* muscles of the rat was measured with a resolution of 100 ms, capturing the recovery process. The curve obtained from measuring the integral of the absolute value of EMG responses to stimuli during and after block was fitted to a sigmoid which enables the identification of 3 regions of recovery. The initial region is of low EMG activity and corresponds to the portion of the stimulus train delivered during block. The second region captures the recovery of the muscle from block, and the third corresponds to the steady state of muscle activation after the recovery period.

Results for both muscles were collated to provide trends using box plots observed in the data. Showing that as the blocking signal amplitude and frequency are increased, the recovery time increases and the normalised EMG response after recovery decreases, indicating that the muscle and thereby the nerve has more trouble recovering within the trial’s time frame. Large variance was seen in the results, motivating further work and refinement of the protocol to validate the trends observed in the data.

## Acknowledgement

The authors gratefully acknowledge Dr. Ian Williams of the Centre for Bio-inspired Technology, Imperial College London, SW7 2AZ UK for his assistance in reviewing this manuscript. This work is supported by the Engineering & Physical Sciences Research Council (EPSRC), UK (SenseBack project, grant EP/M025977/1). Additionally, AR is supported through the EPSRC Centre for Doctoral Training (CDT) in High Performance Embedded & Distributed Systems (HiPEDS, EP/L016796/1), EB and KN through EPSRC grants: EP/M025594/1 and EP/N023080/1.

## Open Access Statement

Upon acceptance, this work will be made Open Access, under the CCBY Creative Commons license. The EMG data supporting this publication and additional metadata will be made freely available via the Newcastle University Research Data Service.

